## Supplementary Figures for "The combined action of CTCF and its testis-specific paralog BORIS is essential for spermatogenesis"

Rivero-Hinojosa et al.

### Supplementary Figures 1-15

**Supplementary Figure 1**

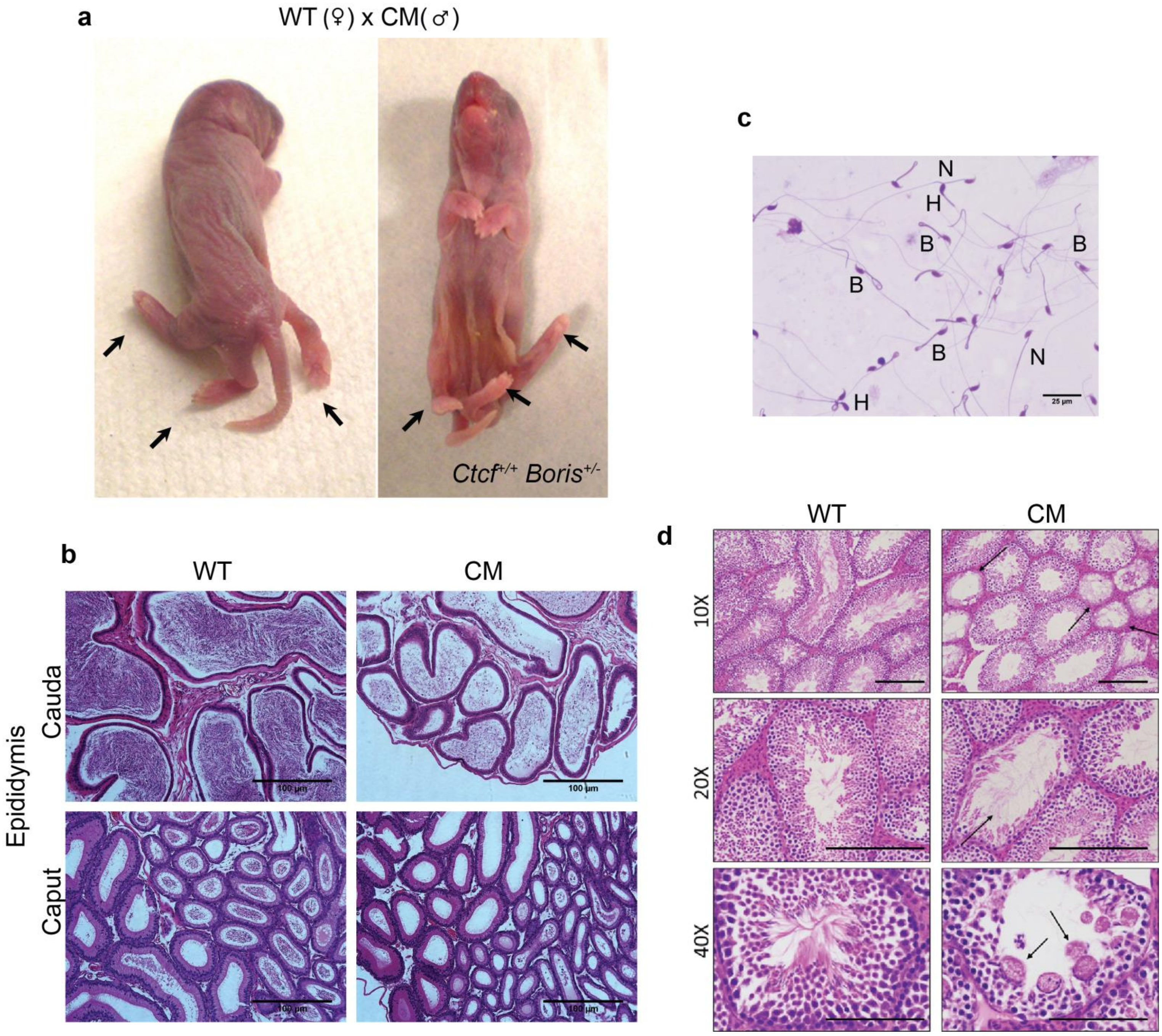

**Supplementary Figure 1. CM mice show spermatogenic defects. a** Pictures of stillborn pup with an extra rear extremity (fifth leg). **b** Histological H&E staining of WT and CM caput and cauda epididymis. Scale bars: 100μm. **c** Visualization by light microscopy of spermatozoa from CM mice. The image demonstrates the cumulative spermatozoa defects in sterile CM mice, including not only a dramatic loss of the total sperm counts but also severe deformations, with tails bent 180˚ through the cytoplasmic droplet indicated with the letter B and abnormal rounded head shape with the letter H. Whereas a small residual number of normal sperms was observed, indicated with the letter N. Scale bars: 25μm. **d** Hematoxylin and eosin staining of testes prepared from 3-month-old WT and CM males. Upper panel: Histological H&E staining of WT and CM tubules at 10X magnification. Black arrows show the empty tubules with a smaller number of meiotic and post meiotic cells in CM tubules. Middle panel: Histological H&E staining of WT and CM tubules at 20X magnification. Black arrows show CM tubule without post meiotic cells. Lower panel: Histological H&E staining of WT and CM tubules at 40X magnification. Black arrows show multinucleated giant cells in CM tubule. Scale Bars: 200 μm (upper and middle panels), 100 μm (lower panels). For **b**, **c** and **d** 7 animals for each type of mice were analyzed with similar results.

**Supplementary Figure 2**

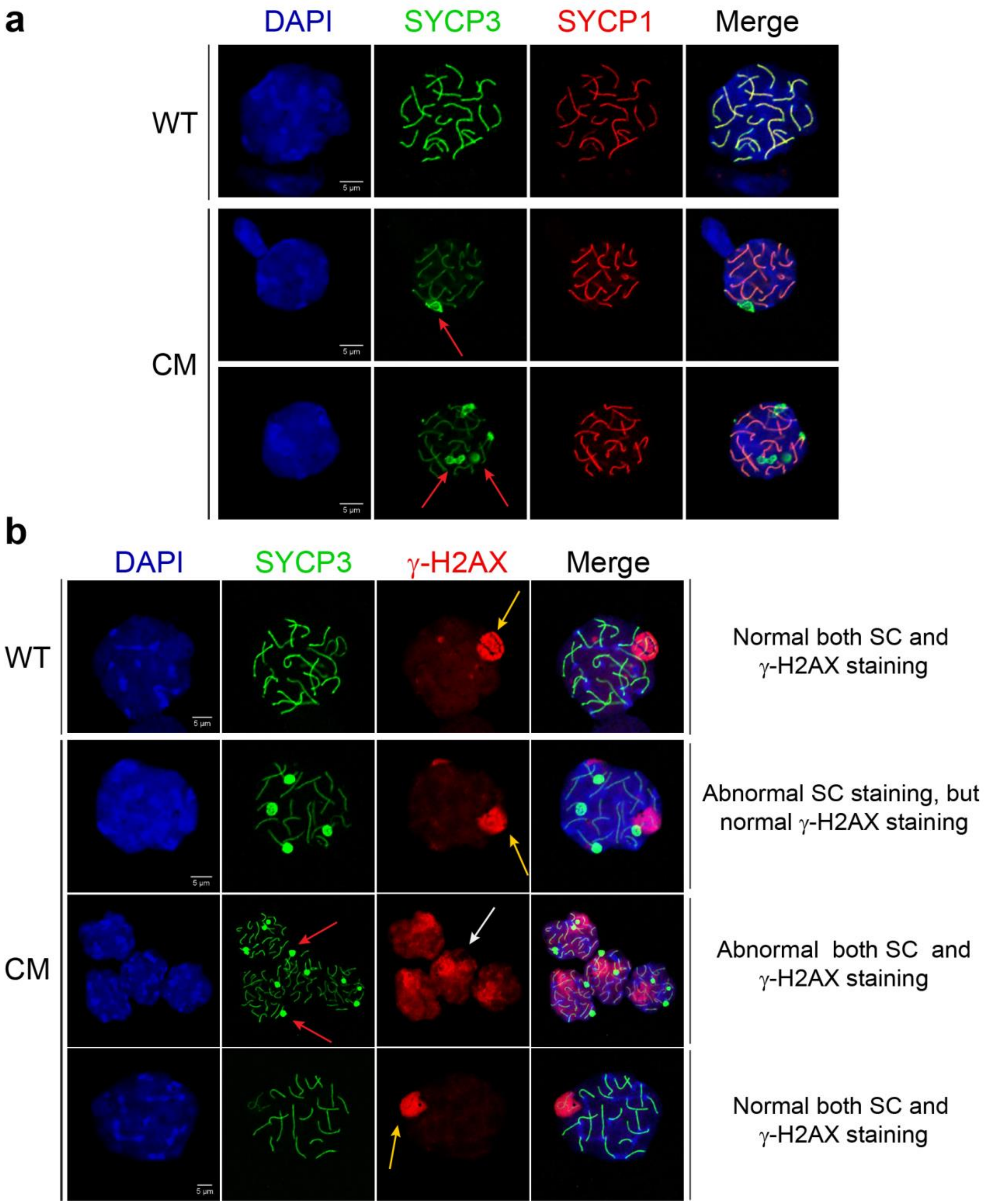

**Supplementary Figure 2. CM mice show defects in Synaptonemal Complex (SC) formation. a** Pachytene spermatocytes prepared from WT and CM mouse testes were fixed and analyzed by immunofluorescence microscopy. Spermatocyte spread preparations were stained with anti-SYCP3 (green) and SYCP1 (red) antibodies for visualization of the lateral and central synaptonemal complex components, respectively. The red arrows show the abnormal SYCP3 staining. **b** Spermatocyte spread preparations were stained with anti-SYCP3 (green) and gamma-H2AX (red) antibodies. The unsynapsed chromosomal regions are shown by red arrows. The extension of gamma-H2AX staining beyond the sex vesicle in CM spermatocytes is shown by white arrow, while normal gamma-H2Ax staining is shown by yellow arrows. Two different abnormal formations of the synaptonemal complex are observed in CM compared to WT spermatocytes (summarized on the right). Five animals from each type of mice were analysed with similar results. Scale bars: 5 μm**.**

**Supplementary Figure 3**

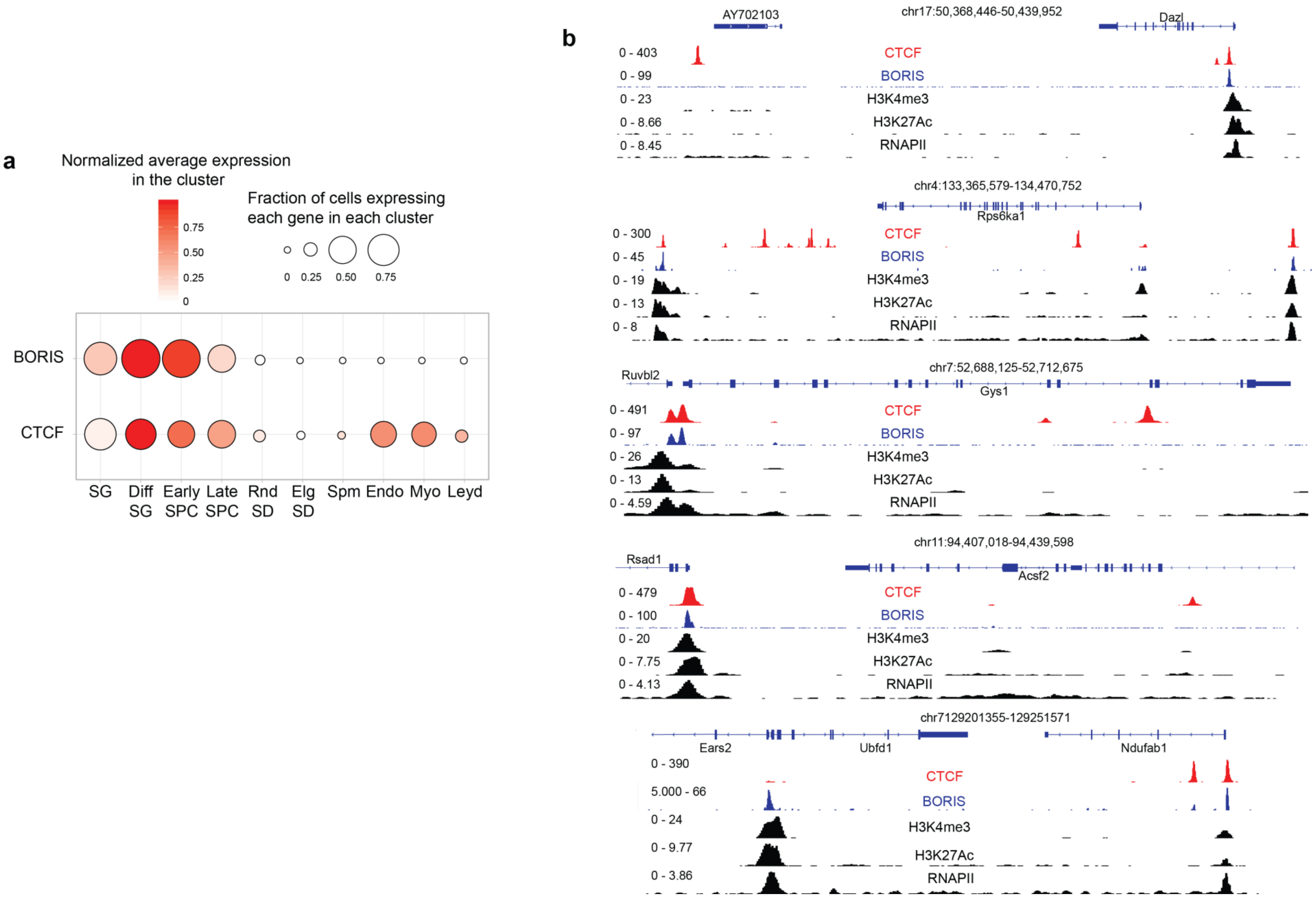

**Supplementary Figure 3**. **Genome wide mapping of CTCF and BORIS binding regions in mouse germ cells by ChIP-seq**. **a** Human single-cell RNA-seq analysis of *CTCF* and *BORIS* expression at the major stages of spermatogenesis based on Guo et al., 2018 (SG: Spermatogonia, SPC: Spermatocytes, SD: Spermatids). **b** Gene tracks for the distribution of CTCF (red) and BORIS (blue) bound regions in mouse male germ cells in correlation with RNA Polymerase II (RNAPII) and active histone marks from ENCODE (H3K4me3 (active promoters) and H3K27ac (active enhancers)).

**Supplementary Figure 4**

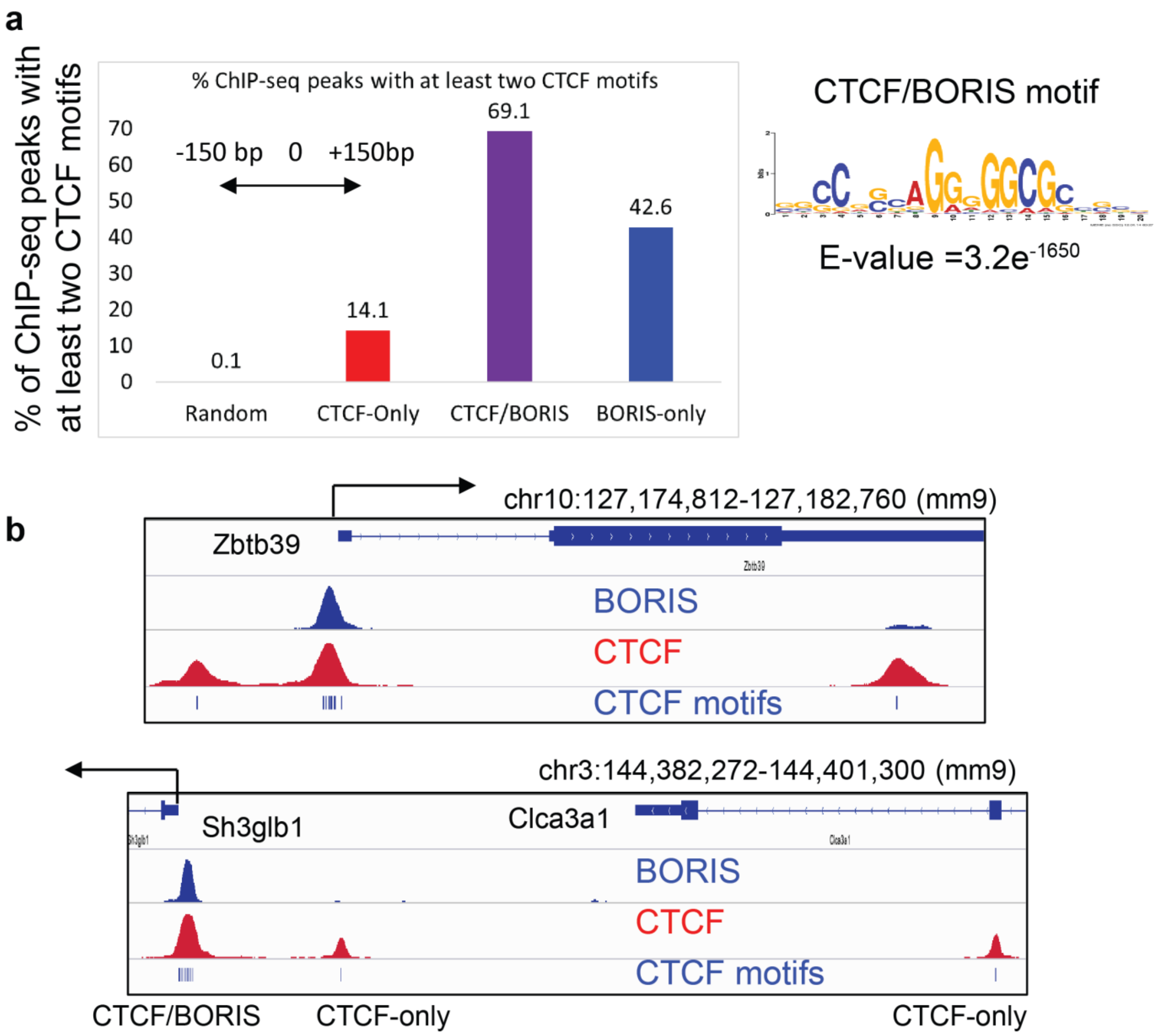

**Supplementary Figure 4. CTCF/BORIS binding regions enclose at least two closely located CTCF binding sites in contrast to CTCF-only binding sites. a** Bar chart shows the percentage of random, CTCF-only, CTCF/BORIS and BORIS-only peaks with two or more CTCF/BORIS motifs. The top 1000 CTCF-only, CTCF/BORIS and BORIS-only binding regions (the regions with the highest tag enrichment) were selected for the analysis. The presence of a CTCF motif was calculated by FIMO (MEME suite) in the sequence extended 150 bp upstream and downstream of the summit of either CTCF (CTCF-only and CTCF/BORIS) or BORIS (BORIS-only) peaks. Each CTCF motif (right panel) occurrence has a p value < 0.0001. **b** Gene tracks show the enrichment of CTCF motifs under CTCF/BORIS binding regions in contrast to CTCF-only binding regions. CTCF and BORIS binding sites in mouse testes are highlighted by red and blue colors, respectively. CTCF motifs are shown by blue bars.

**Supplementary Figure 5**

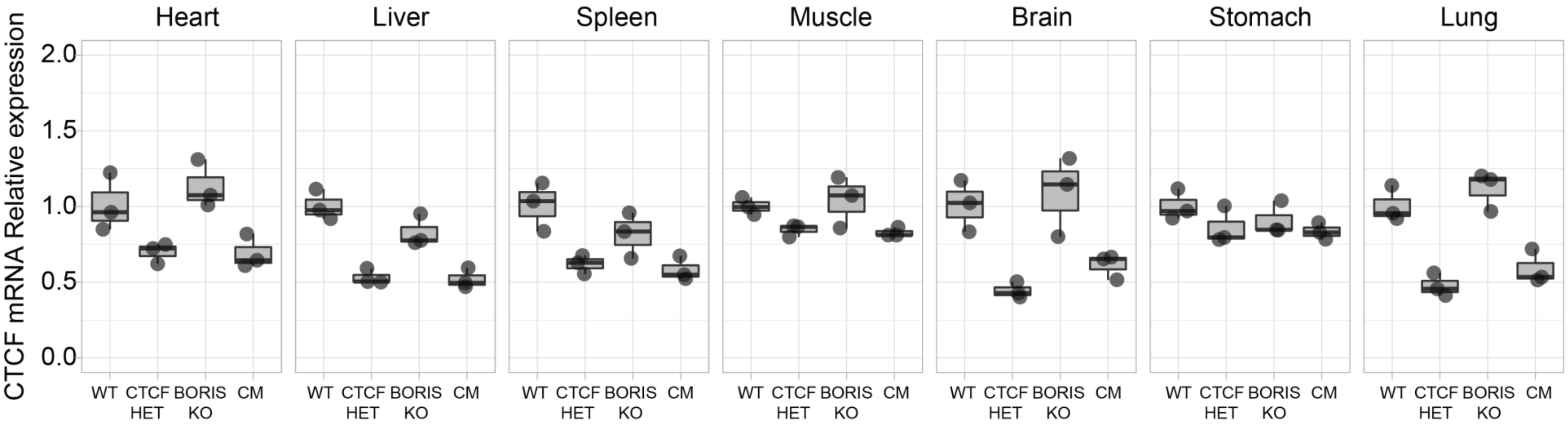

**Supplementary Figure 5. *Boris* knockout has no effect on *Ctcf* expression in somatic tissues**. Ctcf expression by real time PCR in heart, liver, spleen, muscle, brain, stomach, and lung in the four types of mice. The relative *Ctcf* expression level was normalized to *Gapdh* level (n=3). In the Box plots, the lower and upper hinges correspond to the first and third quartiles, the middle line indicates the median. The upper whisker extends from the hinge to the largest value no further than 1.5 times of the inter-quartile range (IQR). The lower whisker extends from the hinge to the smallest value at most 1.5 times of the IQR.

**Supplementary Figure 6**

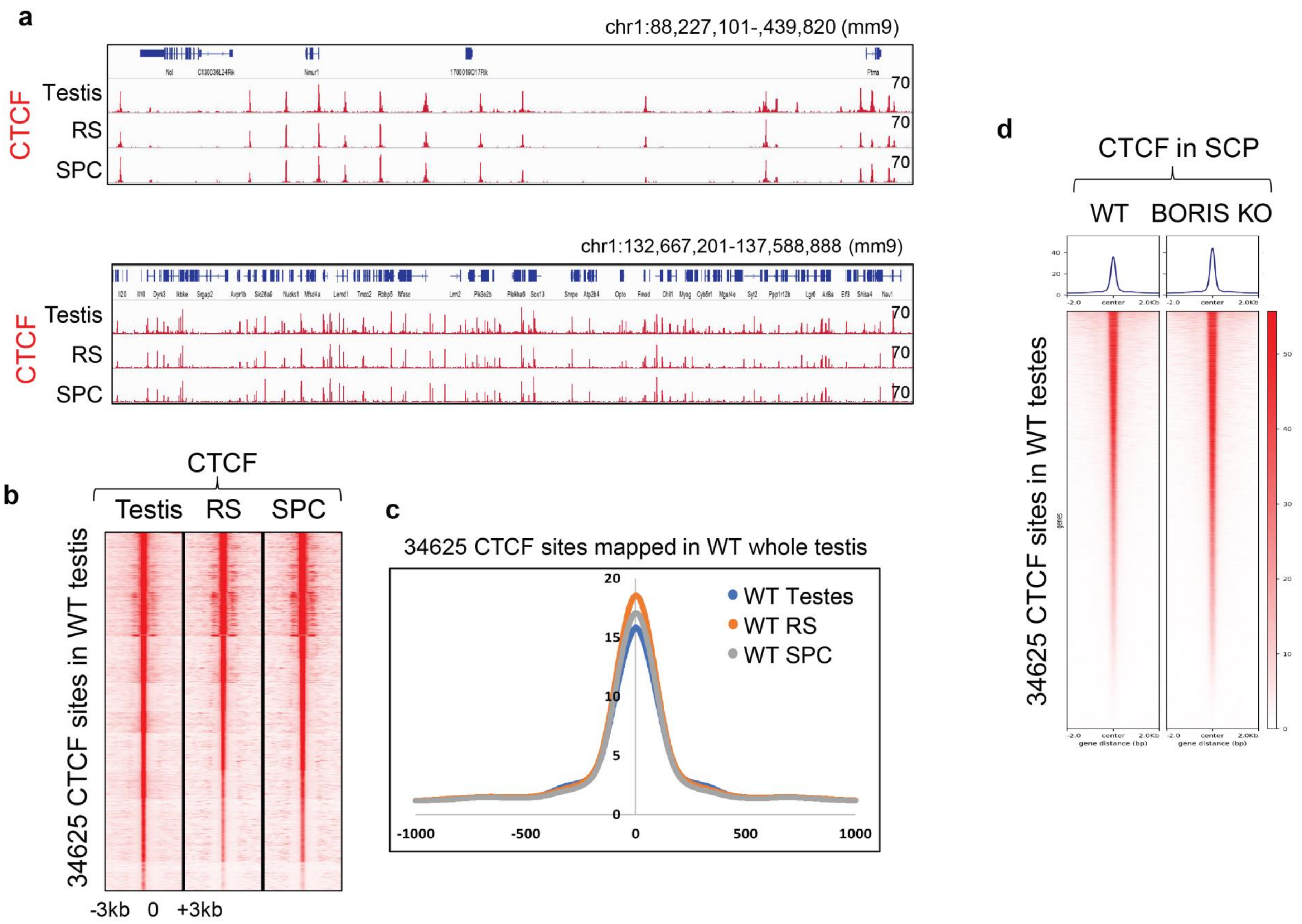

**Supplementary Figure 6. CTCF occupancy mapped in wild type mouse whole testes reflects CTCF occupancy mapped in purified round spermatids and spermatocytes. a** CTCF ChIP-seq tracks represent the similarity of CTCF occupancy in wild type whole testis (Testis), round spermatids (RS) and spermatocytes (SPC). ChIP-seq tag density was normalized to the lowest number of ChIP-seq reads mapped in round spermatids. **b** Heatmap depicts CTCF (red) occupancy in wild type whole testis, RS and SPC at the 34625 CTCF binding sites mapped in whole testis. The tag density of CTCF ChIP-seq data was collected within a 6-kb window around the summit of CTCF peaks mapped in whole mouse testis. The tag density was subjected to k-means enrichment linear with 10 clusters expected. **c** Average tag density of CTCF occupancy mapped in WT whole testes (WT Testes), round spermatids (WT RS) and spermatocytes (WT SPC). The tag density was normalized to the number of mapped reads. **d** Heatmap depicts CTCF (red) occupancy in wild type and BORIS KO SPC at the 34625 CTCF binding sites mapped in whole testes. The tag density of CTCF ChIP-seq data was collected within a 4-kb window around the summit of CTCF peaks mapped in whole mouse testes. The tag density was subjected to k-means enrichment linear.

**Supplementary Figure 7**

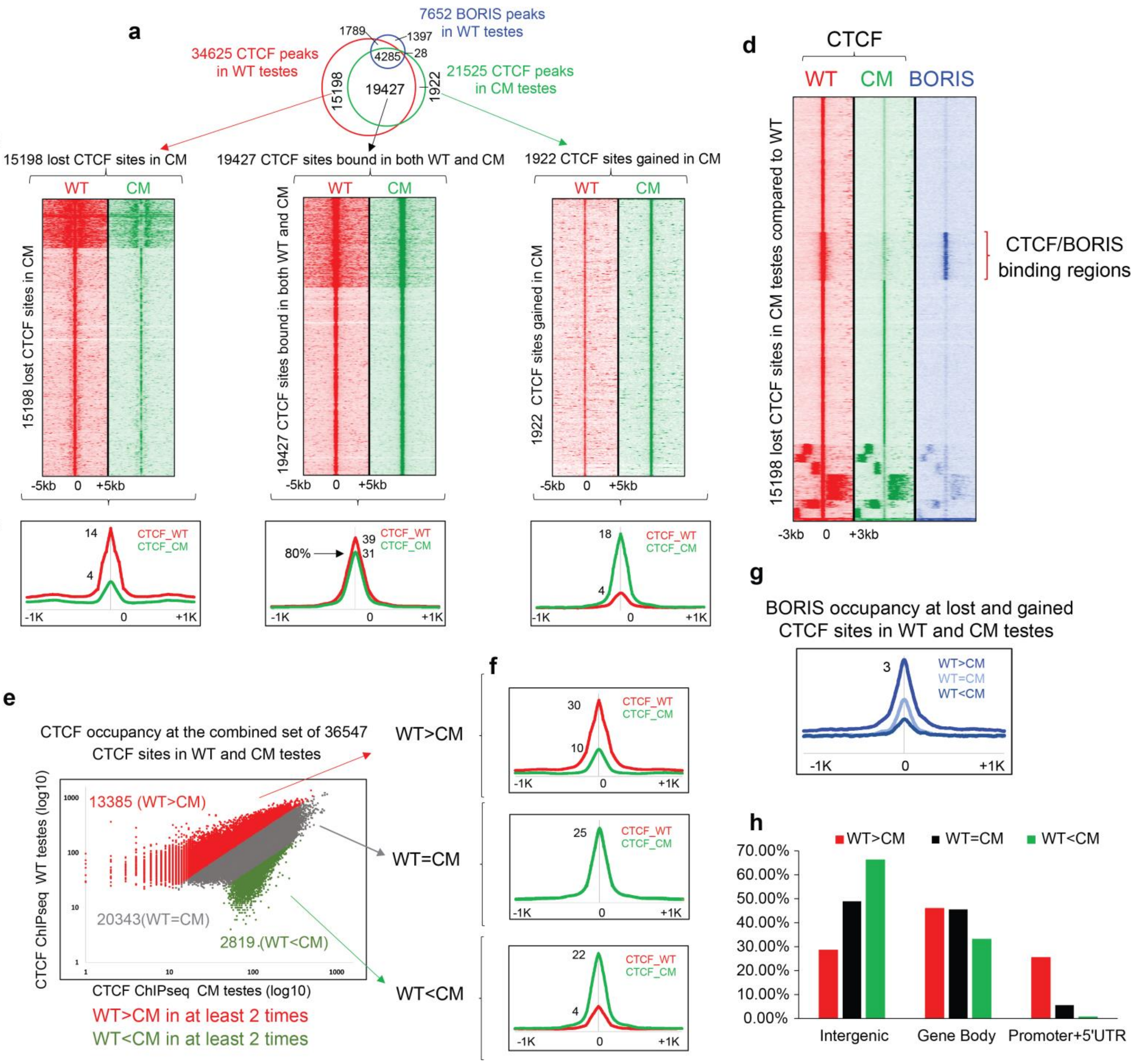

**Supplementary Figure 7. Loss of genome wide CTCF occupancy in CM compared to WT testes.** **a** Venn diagram shows an overlapping of CTCF ChIP-seq peaks mapped in WT (red) and CM (green) testes, and BORIS ChIP-seq peaks mapped in WT (blue) testes. **b** Heatmap of CTCF tag density in WT (red) and CM (green) testes at CTCF binding regions from overlap in panel A. The connections between panels **a** and **b** are shown by arrows. **c** Average tag density of CTCF occupancy in WT (red) and CM (green) testes at the CTCF binding regions from panel **a**. **d** Heatmap of CTCF tag density in WT (red) and CM (green) testes and BORIS tag density (blue) mapped in WT testes generated by K-means clustering of ChIP-seq data along the 15198 lost CTCF binding regions. The cluster of CTCF binding sites lost in CM testes compared to WT is associated with CTCF/BORIS binding regions in WT testes (shown by red bracket). **e** Scatter plot of CTCF occupancy in WT and CM testes at the combined set of 36547 CTCF binding regions. Normalized ChIP-seq tag density was calculated at each CTCF binding region (500 bp upstream and downstream of the summit of CTCF ChIP-seq peak). **f** Average tag density of CTCF occupancy in WT (red) and CM (green) testes at the CTCF binding regions from panel **e**. WT=CM CTCF binding regions were selected with the tag density difference of no more than 1.2-fold between WT and CM testes (total 7051 CTCF binding regions). **g** Average tag density of BORIS occupancy at the CTCF binding regions from panel **f**. **h** Genomic distribution of differentially bound CTCF regions from panel **e** at promoters +5'UTR, gene body, and intergenic regions. CTCF regions differentially bound in WT are associated with promoter regions.

**Supplementary Figure 8**

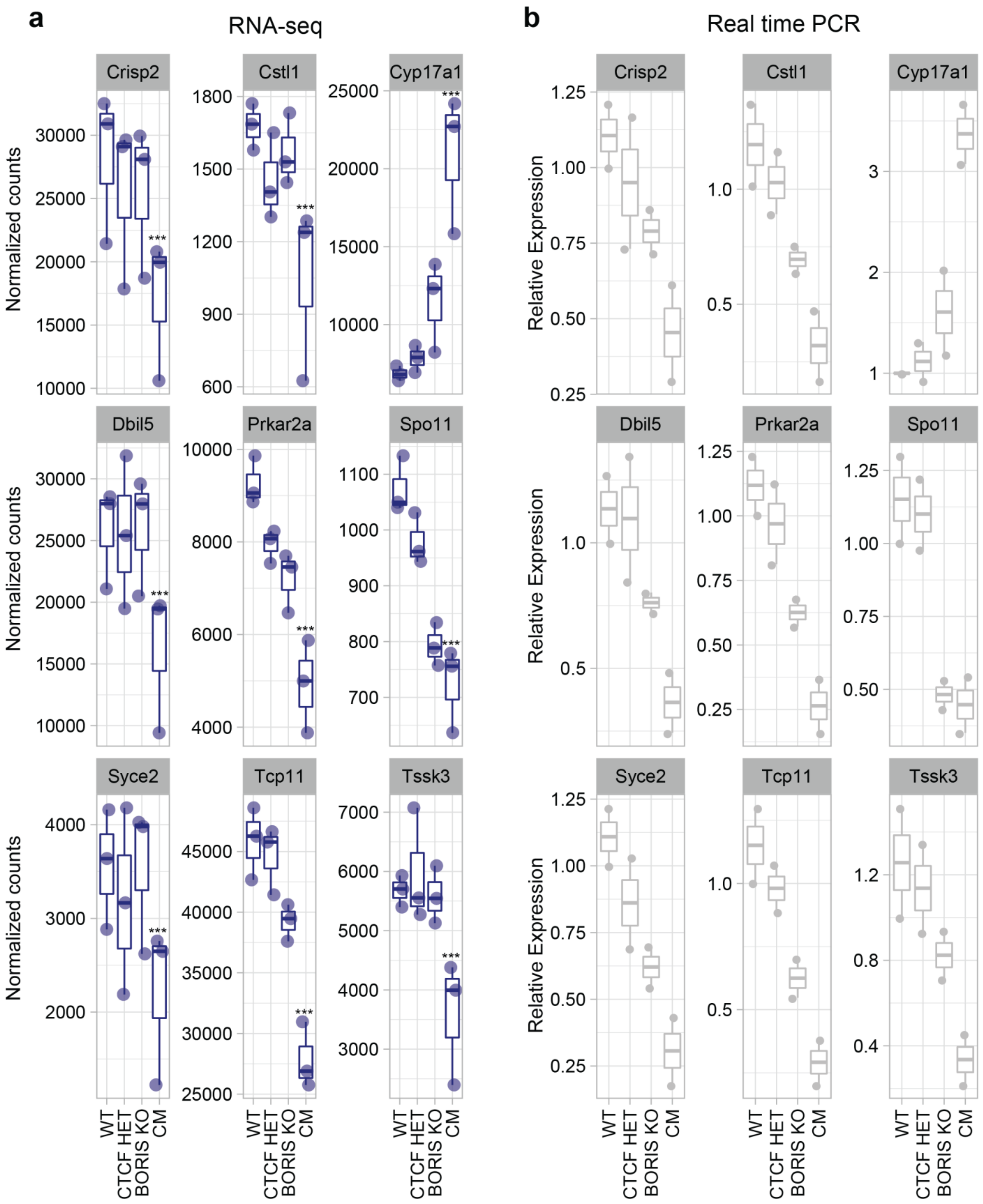

**Supplementary Figure 8. Real time quantitative PCR verification of RNAseq results.** Validation of the RNA-seq results by RT-qPCR of 9 differentially expressed genes. **a** Blue: normalized count values for each gene derived by RNA-seq analysis for each type of mice, p-values were calculated using DESeq2, where the p-values attained by the Wald's test were corrected for multiple testing using the Benjamini and Hochberg method, ***p-values CM versus WT (n=3): *Crisp2*=0.0123, *Cstl1*=0.0187, *Cyp17a1*=1.20e-16, *Dbil5*=0.0256, *Prkar2a*=7.55e-06, *Spo11*=0.0004, *Syce2*=0.0324,*Tcp11*=7.48e-06, *Tssk3*= 0.0081. **b** Grey: RT-qPCR expression for each gene. The expression level of each gene was normalized to Gapdh in RT-qPCR experiments (n=2). In the Box plots, the lower and upper hinges correspond to the first and third quartiles, the middle line indicates the median. The upper whisker extends from the hinge to the largest value no further than 1.5 times of the inter-quartile range (IQR). The lower whisker extends from the hinge to the smallest value at most 1.5 times of the IQR.

**Supplementary Figure 9**

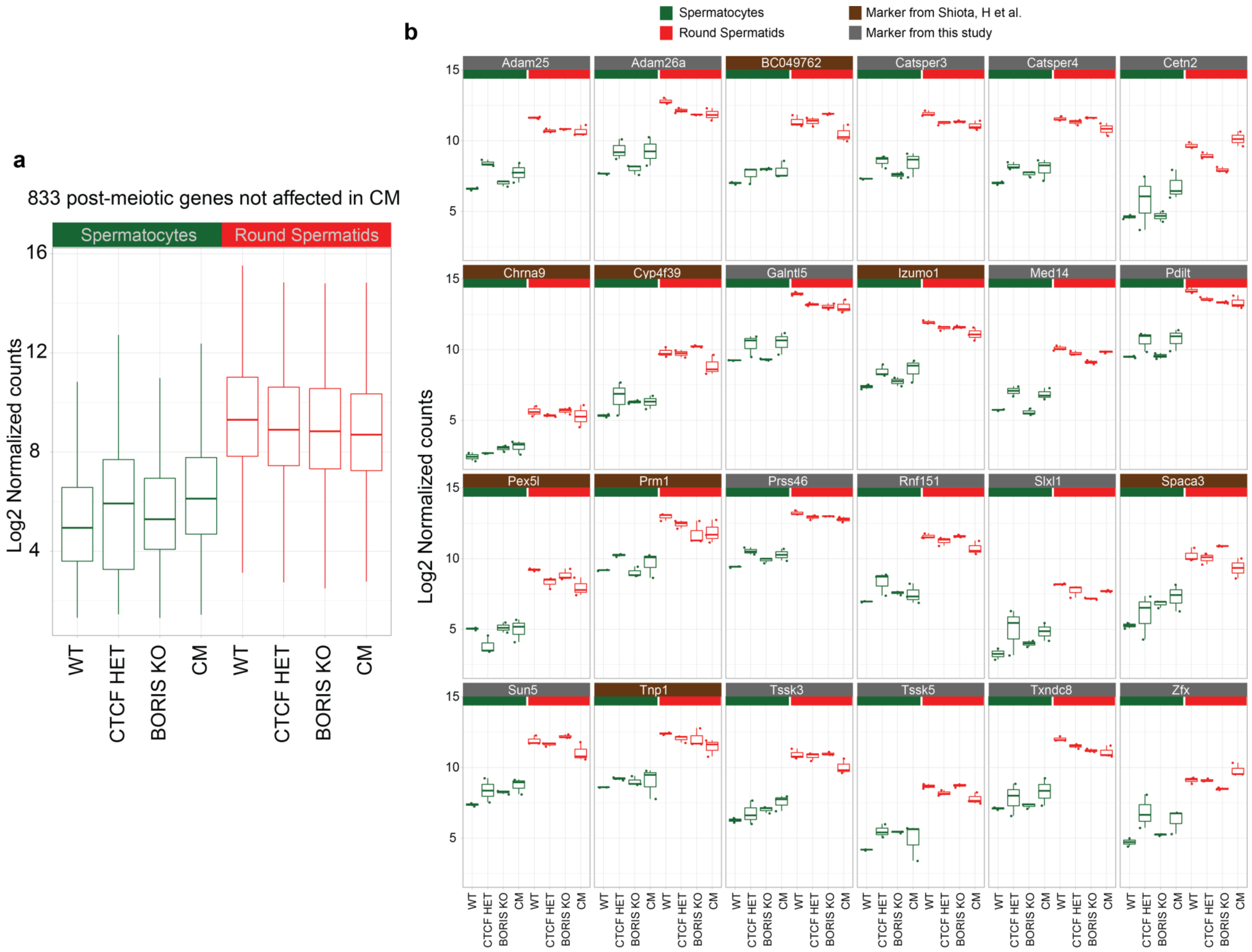

**Supplementary Figure 9. Expression profile of genes up-regulated in post-meiotic cells in the four types of mice**. **a** Quality control of WT, CTCF HET, BORIS KO and CM spermatocyte (green) and round spermatid (red) fractions. In order to confirm that post-meiotic WT and CM populations are comparable, the global expression of 833 genes that are activated in post-meiotic cells compared to spermatocytes, was monitored (n=833). Differentially expressed genes in CM whole testis were excluded from this list of genes. **b** Expression of 24 individual round spermatid marker genes within the genes which expression is globally shown in (a), (n=3). Sixteen markers were selected by comparing WT round spermatids to WT spermatocytes. Eight markers where selected from a similar analysis by Shiota et al., 2018, Cell Reports 24, 3477–3487. In the Box plots, the lower and upper hinges correspond to the first and third quartiles, the middle line indicates the median. The upper whisker extends from the hinge to the largest value no further than 1.5 times of the inter-quartile range (IQR). The lower whisker extends from the hinge to the smallest value at most 1.5 times of the IQR.

**Supplementary Figure 10**

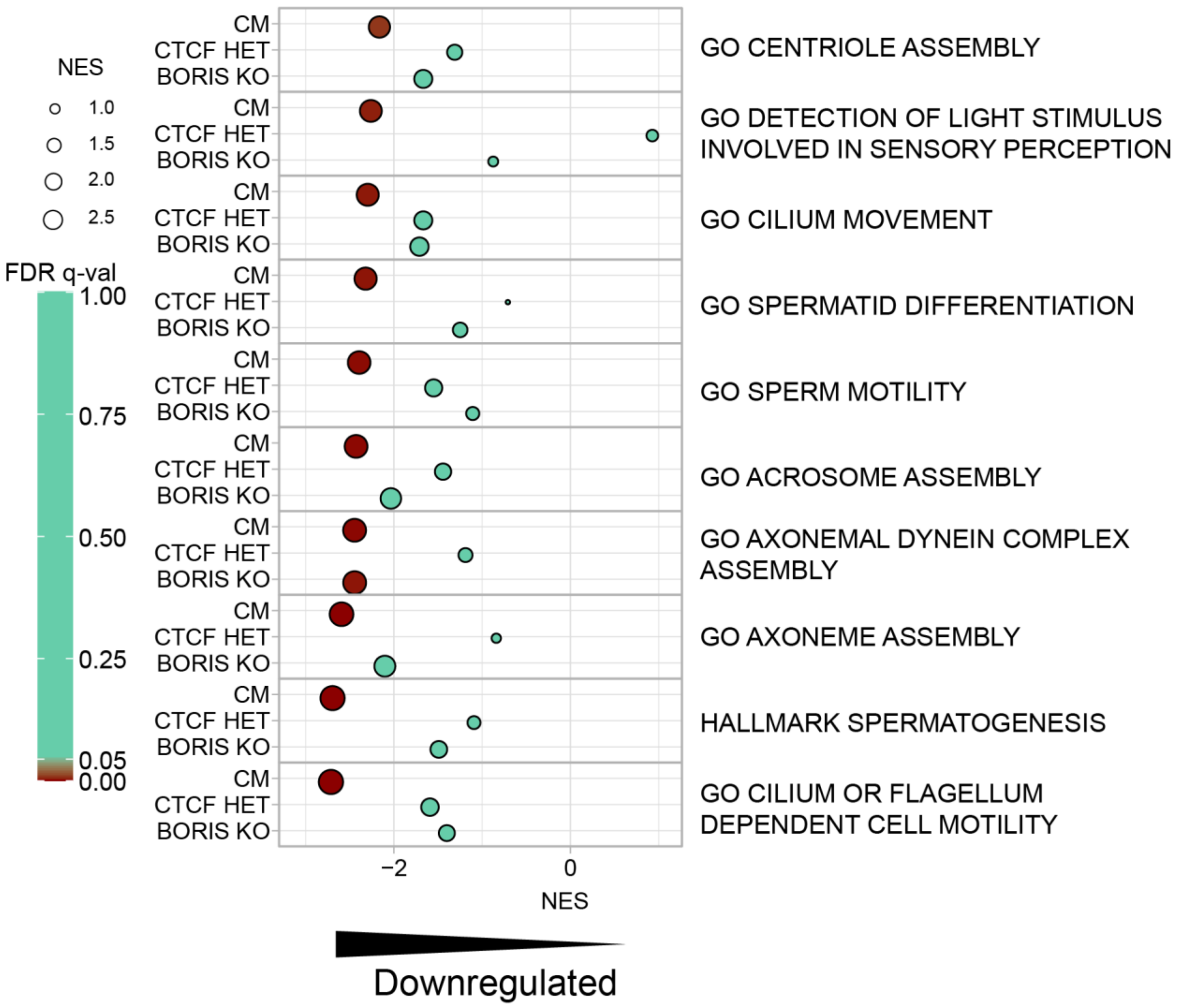

**Supplementary Figure 10. Synergistic effect of CTCF and BORIS depletion on spermatogenesis related pathways.** Gene set enrichment analysis (GSEA) in CM, CTCF HET and BORIS KO whole testes compared to WT whole testes. The normalized enrichment score (NES) for the top 10 significant gene sets downregulated in CM whole testes are shown for CM versus WT, CTCF HET versus WT and BORIS KO versus WT comparisons. Spermatogenesis associated pathways are significantly downregulated only in CM testes, the same pathways were not significantly enriched in CTCF HET or BORIS KO testes. GSEA was performed using the GSEA v4.1.0 software and the following Molecular Signatures Database (MSigDB): the Hallmark gene sets (g), the Curated gene sets (C2) and the Ontology gene sets (C5), FDR q-values (FDR q-val) from GSEA are shown, significant gene sets in CM testis in dark red red.

**Supplementary Figure 11**

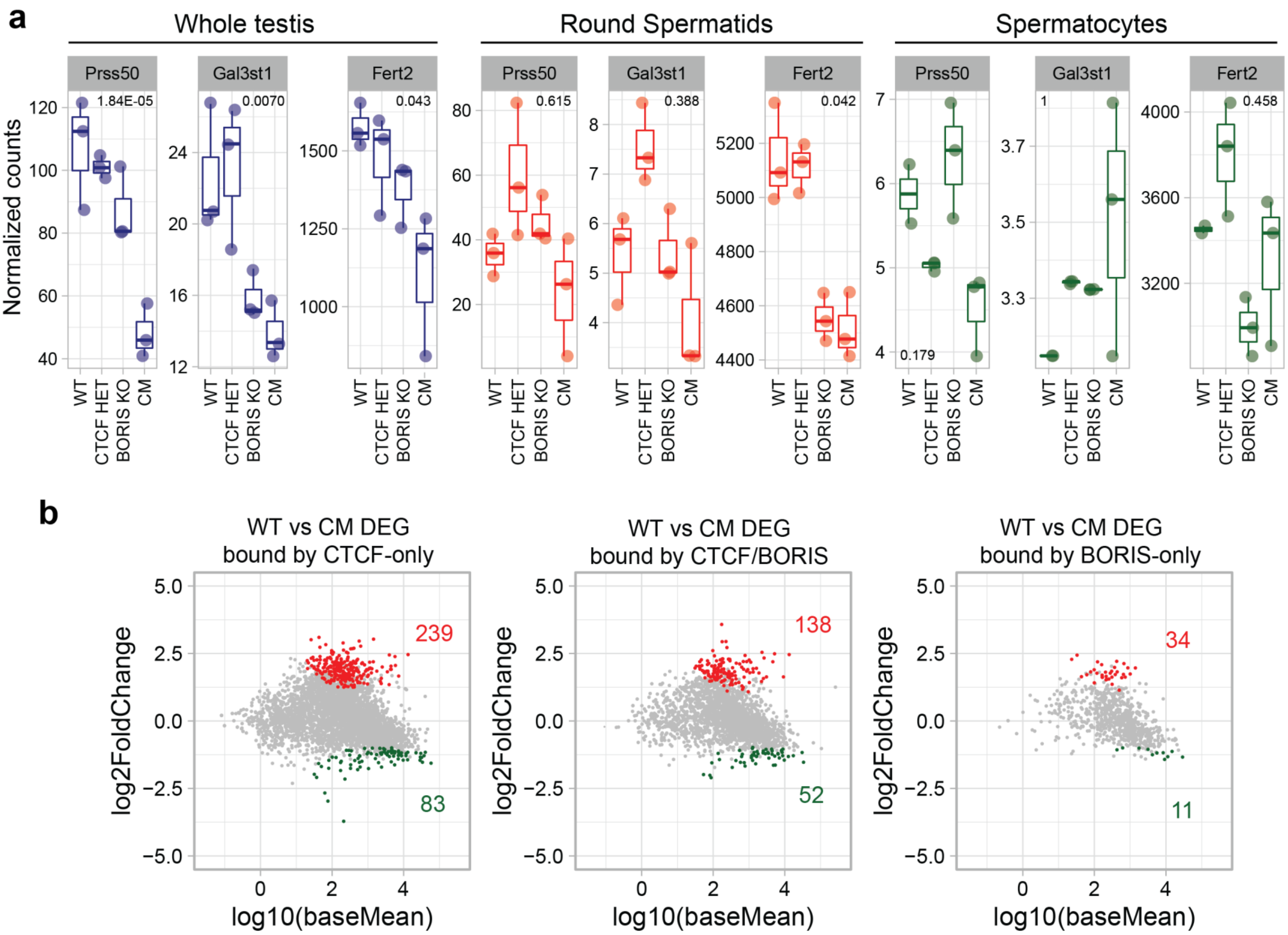

**Supplementary Figure 11. Effect of CTCF and BORIS depletion on spermatogenesis-specific gene expression. a** RNA-seq derived expression levels of different known targets of CTCF and BORIS (*Gal3st1*, *Prss50* and *Fert2*) in whole testis (blue), round spermatids (red) and spermatocytes (green). The y-axis indicates the normalized count values in the RNA-seq analysis. The p-adjusted-values calculated using DESeq2 are indicated in each panel (n=3). DESeq2 calculates the p-values with the Wald's test and corrects for multiple testing using the Benjamini and Hochberg method. The lower and upper hinges correspond to the first and third quartiles, the middle line indicates the median. The upper whisker extends from the hinge to the largest value no further than 1.5 times of the inter-quartile range (IQR). The lower whisker extends from the hinge to the smallest value at most 1.5 times of the IQR. **b** MA plots of RNA-seq for CM mice whole testis on genes with CTCF-only, CTCF/BORIS and BORIS-only sites. The y-axis represents the log2 of fold change compared to WT; the x-axis represents log10 of mean expression. The red and green dots indicate the up- and down-regulated genes, respectively. The gray dots indicate genes without significant changes. The number of differentially expressed genes (DEGs) are highlighted in red (Fold change > 2 and adjusted p-value< 0.001).

**Supplementary Figure 12**

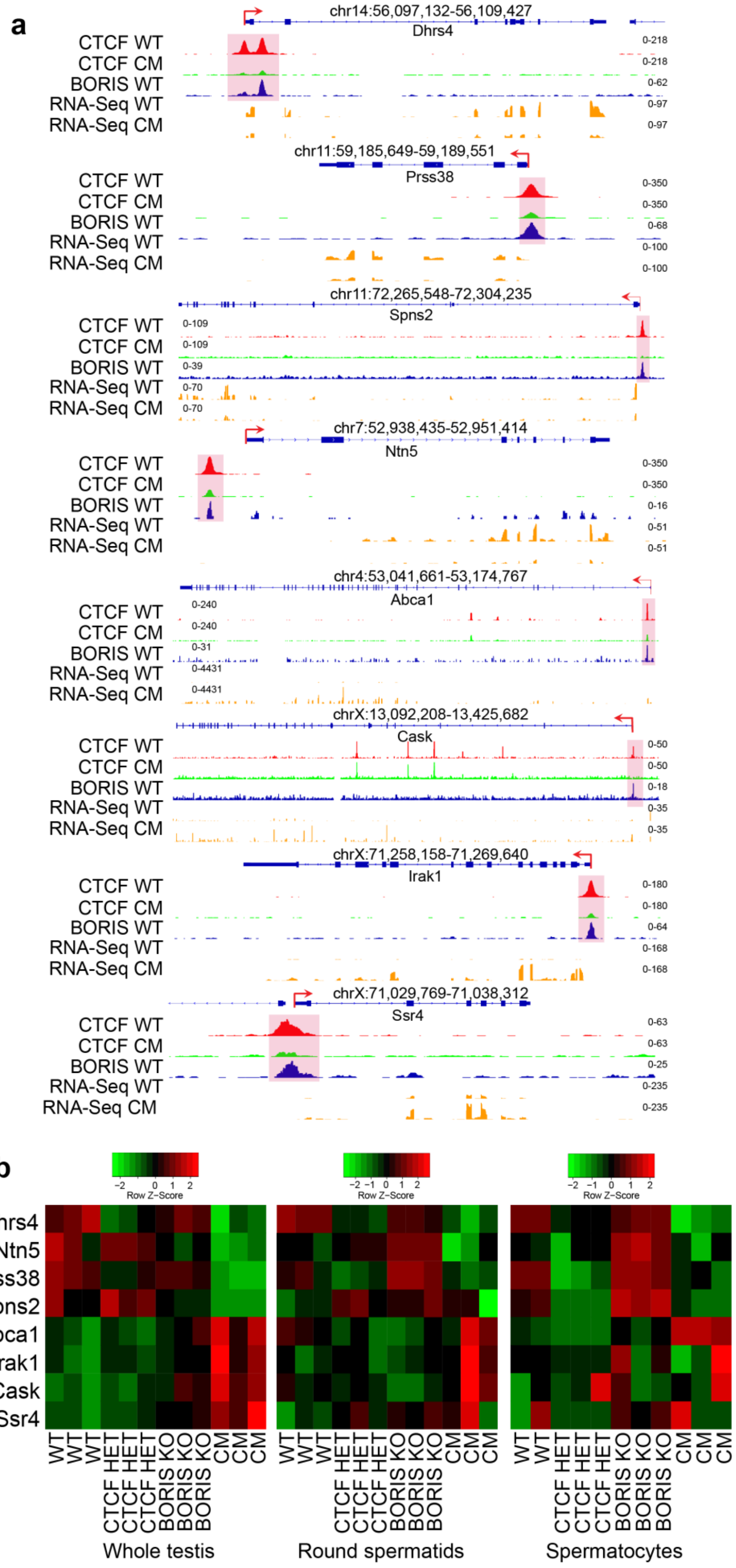

**Supplementary Figure 12. Genome-wide expression analysis in CM germ cells. a** The loss of CTCF binding in CM is associated with differential gene expression. Genomic tracks displaying ChIP-seq for CTCF (WT in red and CM in green) and BORIS (WT in blue), and RNA-seq (yellow) in WT and CM germ cells across the *Dhrs4*, *Prss38*, *Spns2*, *Ntn5*, *Abca1*, *Cask*, *Irak1*, and *Ssr4* loci. **b** Heatmaps showing the change of expression of the same genes shown above in whole testis, round spermatids and spermatocytes of the four types of mice.

**Supplementary Figure 13**

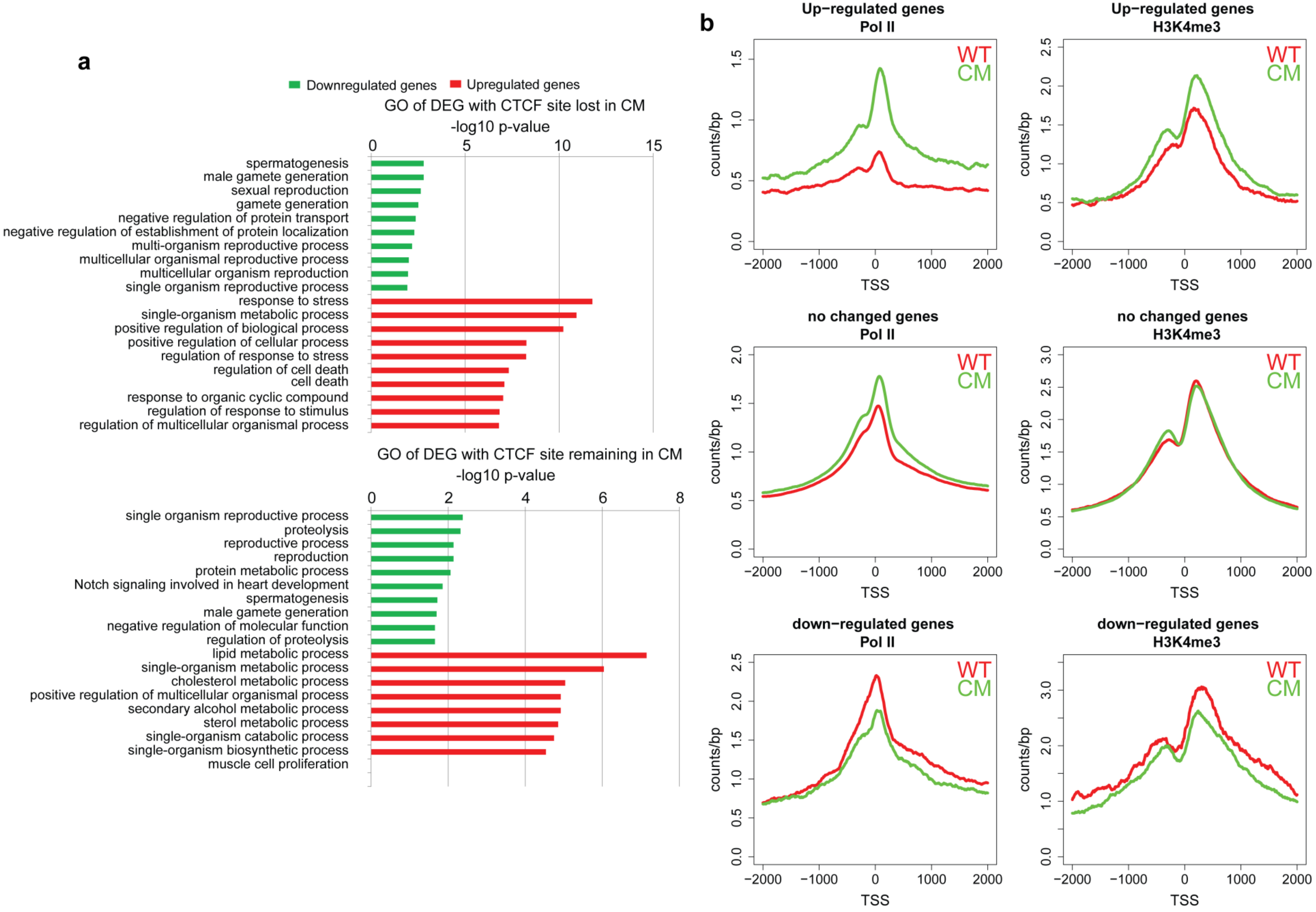

**Supplementary Figure 13. The loss of CTCF binding in CM testis is associated with changes of RNA Pol II and H3K4me3 profiles. a** Gene ontology of differentially expressed genes (dowregulated genes in green and upregulated genes in red) with CTCF site lost (n=327) or remaining (n=148) in its promoter. The analysis was performed in DAVID Bioinformatics Resources 6.8 (https://david.ncifcrf.gov), Fisher's exact p-values from DAVID software are shown. **b** Tag density plots of RNA Polymerase II (Pol II) and H3K4me3 in up-regulated, down-regulated or unchanged genes in WT (red) and CM testis (green).

**Supplementary Figure 14**

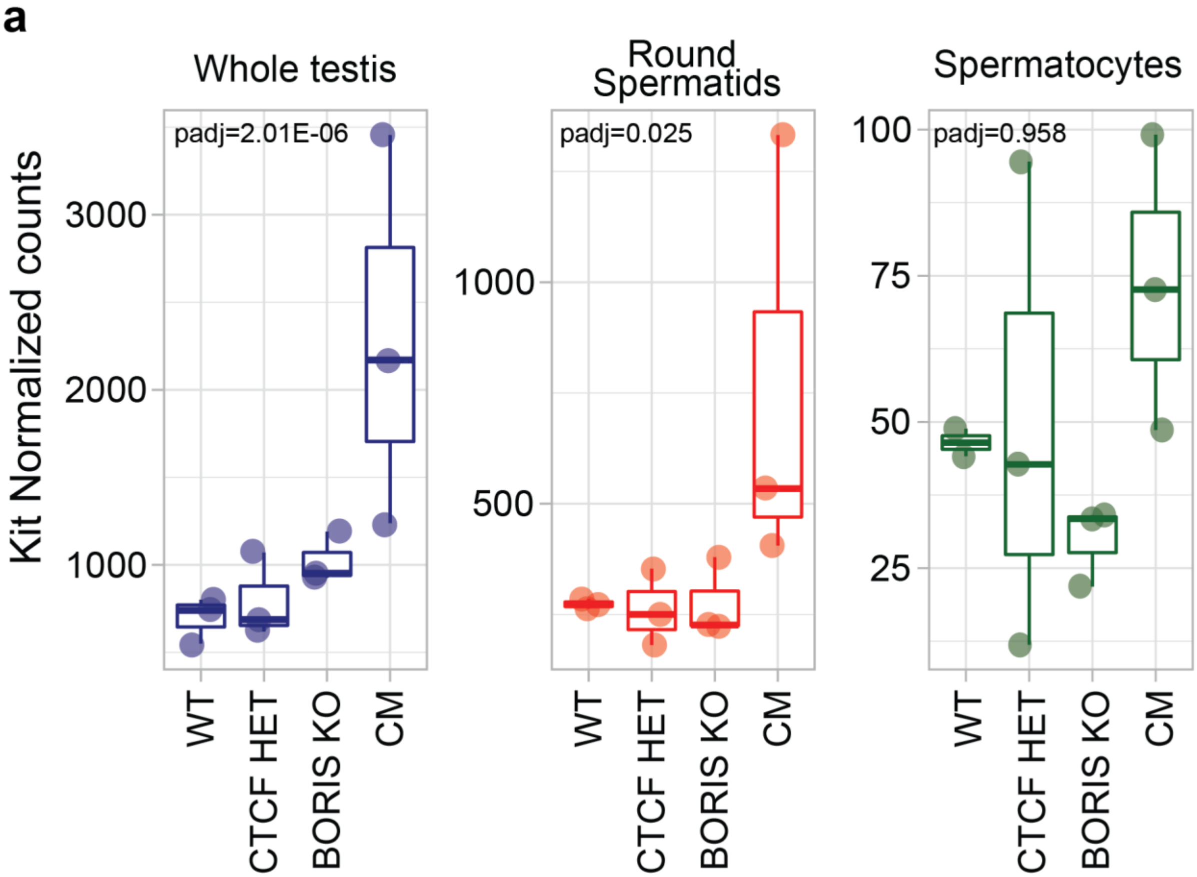


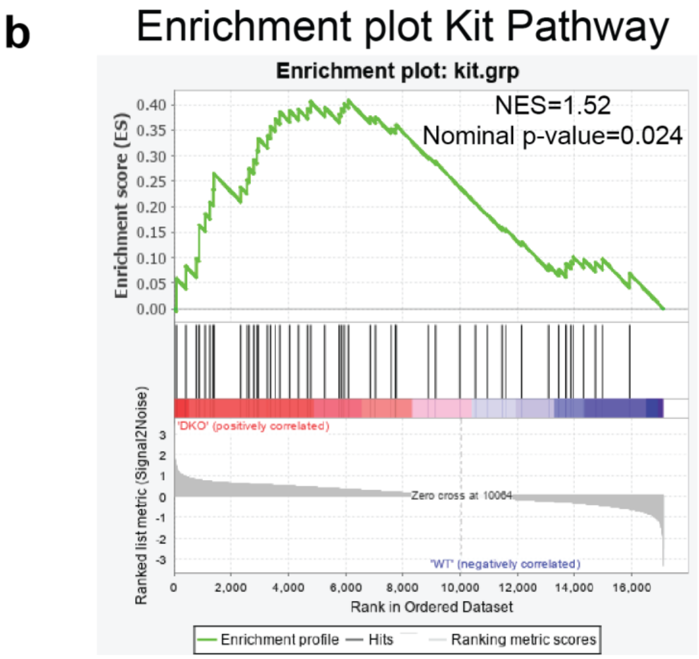

**Supplementary Figure 14. CTCF and BORIS regulate *Kit* expression**. **a** Deletion of both *Boris* and *Ctcf* genes is associated with induction of *Kit* expression. Boxplots representing the expression of *Kit* gene derived from RNA-seq analysis in the four types of mice in whole testis (blue), round spermatids (red) and spermatocytes (green). The p-adjusted-value calculated using DESeq2 is indicated in each panel (n=3). DESeq2 calculates the p-values with the Wald's test and corrects for multiple testing using the Benjamini and Hochberg method. The lower and upper hinges correspond to the first and third quartiles, the middle line indicates the median. The upper whisker extends from the hinge to the largest value no further than 1.5 times of the inter-quartile range (IQR). The lower whisker extends from the hinge to the smallest value at most 1.5 times of the IQR. **b** Gene Set Enrichment Analysis (GSEA) of the KIT pathway performed using the GSEA software. GSEA was performed using the GSEA v4.1.0 software and the following GSEA the Kit Pathway gene set containing 52 genes (https://www.gsea-msigdb.org/gsea/msigdb/cards/PID_KIT_PATHWAY), Nominal p-value and normalized enrichment score (NES) are shown.

**Supplementary Figure 15**

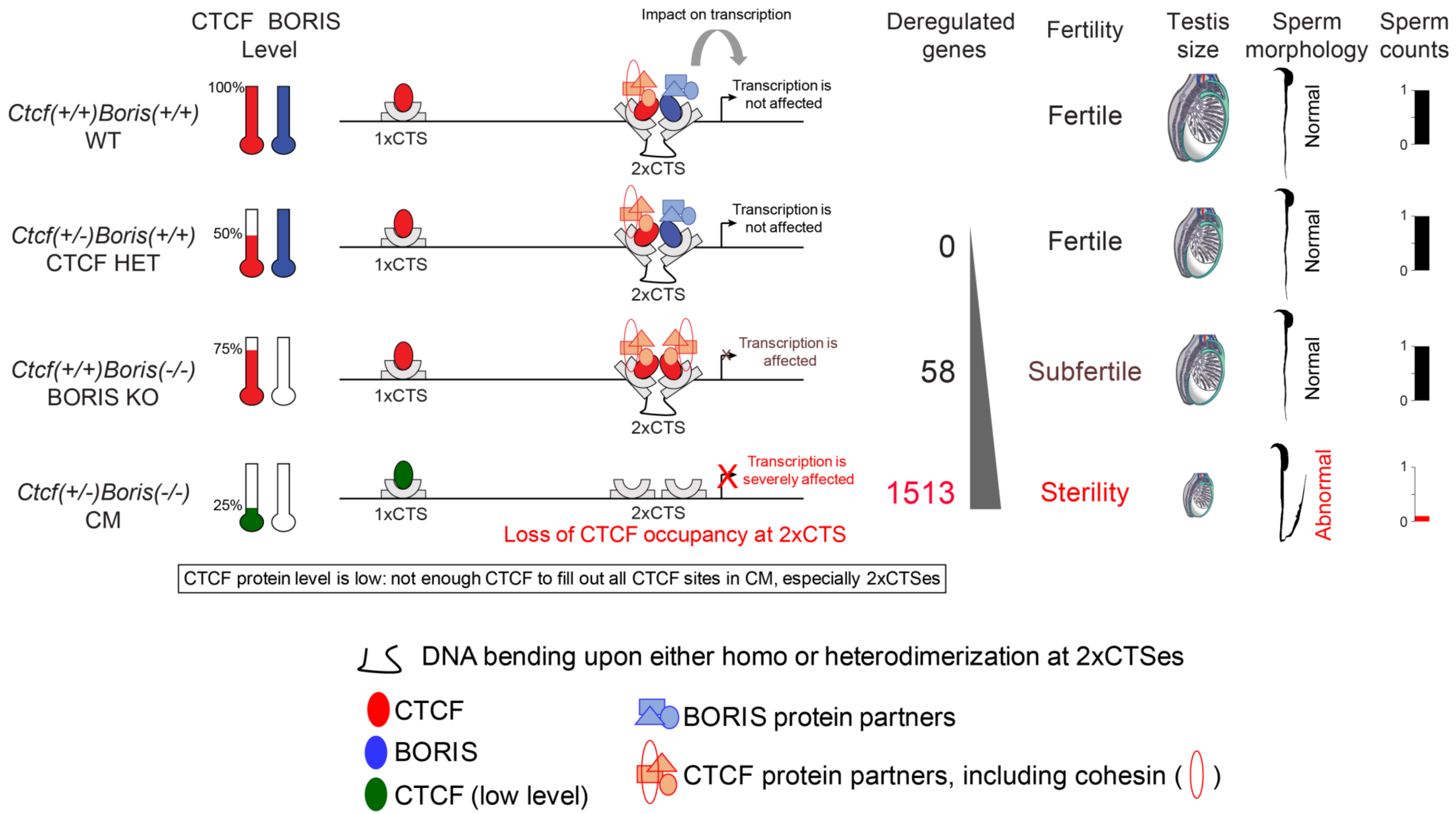

**Supplementary Figure 15. Schematic presentation of the proposed mechanism of how CTCF and BORIS function cooperatively**. A proposed model to explain the non-additive phenotype of CM mice in comparison to WT, CTCF HET and BORIS KO mice. Based on the previously published data, CTCF binds single CTCF binding site (1xCTS) as a monomer, however it forms either homodimer in BORIS-negative somatic cells or heterodimer with BORIS in germ cells at the clustered sites (2xCTS). The 2xCTSes are enriched at promoter regions in contrast to the 1xCTSes, which are largely intergenic. Upon either CTCF homodimerization or CTCF and BORIS heterodimerization at the 2xCTSes, DNA bends to facilitate the creation of higher-order complexes: either CTCF-CTCF-DNA or CTCF-BORIS-DNA. The transcriptional outcome of CTCF and BORIS heterodimerization is different from CTCF homodimerization at the 2xCTSes, as the two paralogous proteins recruit different protein partners due to their distinct N- and C-termini. In BORIS KO germ cells, CTCF-BORIS heterodimers at the 2xCTSes are replaced by CTCF homodimers to partially compensate for the loss of BORIS. However, CTCF homodimers cannot completely substitute for CTCF-BORIS heterodimers, resulting in gene deregulation and pronounced spermatogenesis defects in BORIS KO testes. As CTCF protein level in CM testes is severely decreased, due to *Ctcf* gene haploinsufficiency and *Boris* knockout, that low level of CTCF is not sufficient to form stable homodimers at the 2xCTSes but may be sufficient for CTCF monomers to occupy the 1xCTSes. Upon such a loss of CTCF at the 2xCTSes in CM testis, the transcription of target genes is severely affected, resulting in a complete sterility with reduced testis size (owing to apoptosis), low count of sperm, spermiogenesis abnormalities, and defects/delays in meiotic recombination.
